# Identification of new targets for the diagnosis of cysts (four) and trophozoites (one) of the eye pathogen *Acanthamoeba*

**DOI:** 10.1101/2024.10.16.618517

**Authors:** Bharath Kanakapura Sundararaj, Manish Goyal, John Samuelson

**Author notes:** Made equal contributions (Bharath Kanakapura Sundararaj, Manish Goyal).

## Abstract

*Acanthamoebae*, which are free-living amoebae, cause corneal inflammation (keratitis) and blindness, if not diagnosed and effectively treated. While trophozoites adhere to and damage the cornea, *Acanthamoeba* cysts, the walls of which contain cellulose and have two layers connected by conical ostioles, are the diagnostic form by microscopy of the eye or of corneal scrapings. We recently used structural and experimental methods to characterize cellulose-binding domains of Luke and Leo lectins, which are abundant in the inner layer and ostioles. However, no antibodies have been made to these lectins or to a Jonah lectin and a laccase, which are abundant in the outer layer. Here we used confocal microscopy to show that rabbit antibodies to recombinant Luke, Leo, Jonah, and laccase generally support localizations of GFP-tagged proteins in walls of transfected *Acanthamoebae.* Rabbit antibodies to all four wall proteins efficiently detected calcofluor white-labeled cysts of 10 of 11 *Acanthamoeba* isolates obtained from the ATCC, including five T4 genotypes that cause most cases of keratitis. Laccase shed into the medium during encystation was detected by an enzyme-linked immunoassay. We also used structural and experimental methods to characterize the mannose-binding domain of an *Acanthamoeba* mannose-binding protein and showed that rabbit antibodies to the mannose-binding domain efficiently detected trophozoites of all 11 *Acanthamoeba* isolates. We conclude that four wall proteins are all excellent targets for diagnosing *Acanthamoeba* cysts in the eye or corneal scrapings, while the mannose-binding domain is an excellent target for identifying trophozoites in cultures of corneal scrapings.

**Importance:** Free-living amoeba in the soil or water cause *Acanthamoeba* keratitis, which is diagnosed by identification of cysts by microscopy of the eye or of corneal scrapings, using calcofluor-white that unfortunately cross-reacts with fungi and plants. Alternatively, *Acanthamoeba* infections are diagnosed by identification of trophozoites in cultures of scrapings. Here we showed that rabbit antibodies to four abundant cyst wall proteins (Jonah, Luke, Leo, and laccase) each efficiently detect calcofluor-white-labeled cysts of 10 of 11 *Acanthamoeba* isolates obtained from the ATCC. Further, laccase released into the medium by encysting *Acanthamoebae* was detected by an enzyme-linked immunoassay. We also showed that rabbit antibodies to the mannose-binding domain of the *Acanthamoeba* mannose-binding protein, which mediates adherence of trophozoites to keratinocytes, efficiently identifies trophozoites of all 11 ATCC isolates. In summary, four wall proteins and the ManBD appear to be excellent targets for diagnosis of *Acanthamoeba* cysts and trophozoites, respectively.

## INTRODUCTION

*Acanthamoeba* is an important eye pathogen in the under-resourced countries, where water for handwashing is scarce, and is an emerging pathogen in the US and Europe, where contact lens use is frequent (1–14). Further, the numbers of *Acanthamoebae*, which are free-living in fresh water and soil, are increasing, because of planet-warming (15). Regardless of the place or route of infection, 18S RNA sequences show the T4 genotype, which includes the genome project Neff strain of *Acanthamoeba castellanii* (Ac), most often causes corneal inflammation (keratitis) (16–21). *Acanthamoebae* cause keratitis when trophozoites adhere to corneal cells via a mannose-binding protein (called here AcMBP) to distinguish it from host mannose-binding proteins (22, 23). Monoclonal antibodies have been made to AcMBP, which may be used to identify trophozoites in cultures of corneal smears, and its mannose-binding domain (ManBD) has been suggested but not experimentally proven (24–28). Adherent trophozoites cause corneal ulcers and scars that may lead to blindness if not successfully diagnosed and treated with topical medications, each of which is difficult to perform (29–35).

*Acanthamoeba* cysts, which form when trophozoites are starved of nutrients, have a cellulose- and chitin-rich wall with two layers (ectocyst and endocyst) connected by conical ostioles (36–42). Cyst walls protect *Acanthamoebae* from killing by disinfectants used to clean surfaces, sterilizing agents in contact lens solutions, and alcohols in hand sanitizers (11, 43–47). Cysts are also the diagnostic form in confocal microscopic examination of the eye and in corneal scrapings of eye lesions (13, 48, 49). Calcofluor white (CFW) and wheat germ agglutinin (WGA), which bind to cellulose and chitin, respectively, in the *Acanthamoeba* cyst wall, are suboptimal diagnostic reagents, as they cross-react with walls of plants and fungi that may irritate the cornea or cause keratitis, respectively (43, 50–53).

The primary goal of the present studies is to test whether abundant proteins in the ectocyst layer (Jonah and laccase) or endocyst layer and ostioles (Luke and Leo), which we identified by mass spectrometry of purified walls and localized by GFP-tagging, are good targets for diagnostic anti-cyst rabbit antibodies (rAbs) (38, 40, 54, 55). Jonah lectins contain one or three β-helical folds (BHFs) like those of Antarctic bacteria, while Ac laccases have three copper oxidase (CuRO) domains like those of bacterial spore coat proteins (56–58). Luke lectins contain two or three β-jelly-roll folds (BJRFs) like those of cellulose-binding domains of bacterial and plant endocellulases, while sets of four disulfide knots (4DKs) of Leo are unique to *Acanthamoeba* (59–63). Here we determined the efficiency of rAbs to Jonah-1, laccase-1, Luke-2, and Leo-A to detect CFW-labeled cysts of 11 isolates of *Acanthamoeba* from the American Type Culture Collection (ATCC), including six T4 genotypes that cause most cases of AK in resourced and under-resourced countries (16–21). We also tested the ability of rAbs to the ManBD of AcMBP, which we proved by its ability to bind to mannose-agarose resin and to human corneal limbal epithelial (HCLE) cells, to detect DAPI-labeled trophozoites of the same 11 isolates of *Acanthamoeba* (24–27, 64).

## RESULTS

### With a single caveat, sequence searches with BLASTP and TBLASTN strongly support the four wall proteins chosen for making rAbs for the diagnosis of *Acanthamoeba* cysts in corneal scrapings

Rabbits were immunized with an *E. coli* maltose-binding protein (EcMBP)-fused to the BHF of Jonah-1 (ACA1_164810), two BJRFs of Luke-2 (ACA1_377670), two sets of 4DKs of Leo-A (ACA1_074730), and the first copper oxidase domain (CuRO-1) of an abundant laccase-1 (ACA1_068450), the AlphaFold structures of which are shown in Figs. 1A, 1B, 1G, and 1H (16, 17, 38, 60, 65, 66). See Fig. S1 for their amino acid sequences and that of the ManBD of AcMBP-1, which is described below. Sequence alignments with BLASTP, which did not include the low complexity spacers that are poorly conserved, showed that the N-terminal BJRF of recombinant Luke-2 was duplicated in Neff strain (ACA1_377510), while the N- and C-terminal 4DKs of recombinant Leo-A have a close paralog (ACA1_083920) (68). The other 15 Luke lectins and 13 Leo lectins, as well as the BHFs of seven other Jonah lectins, of the Neff strain of Ac show <60% positional identities, so that rAbs to recombinant Luke, Leo, and Jonah are unlikely to cross-react with paralogous wall proteins. In contrast, TBLASTN searches of 18 other *Acanthamoeba* genomes in AmoebaDB, the proteins of which have not been predicted, showed the vast majority have >85% positional identity with recombinant Luke, Leo, and Jonah and so are very likely to be detected by rAbs to each protein (16, 17, 36, 67).

**Fig. 1.**
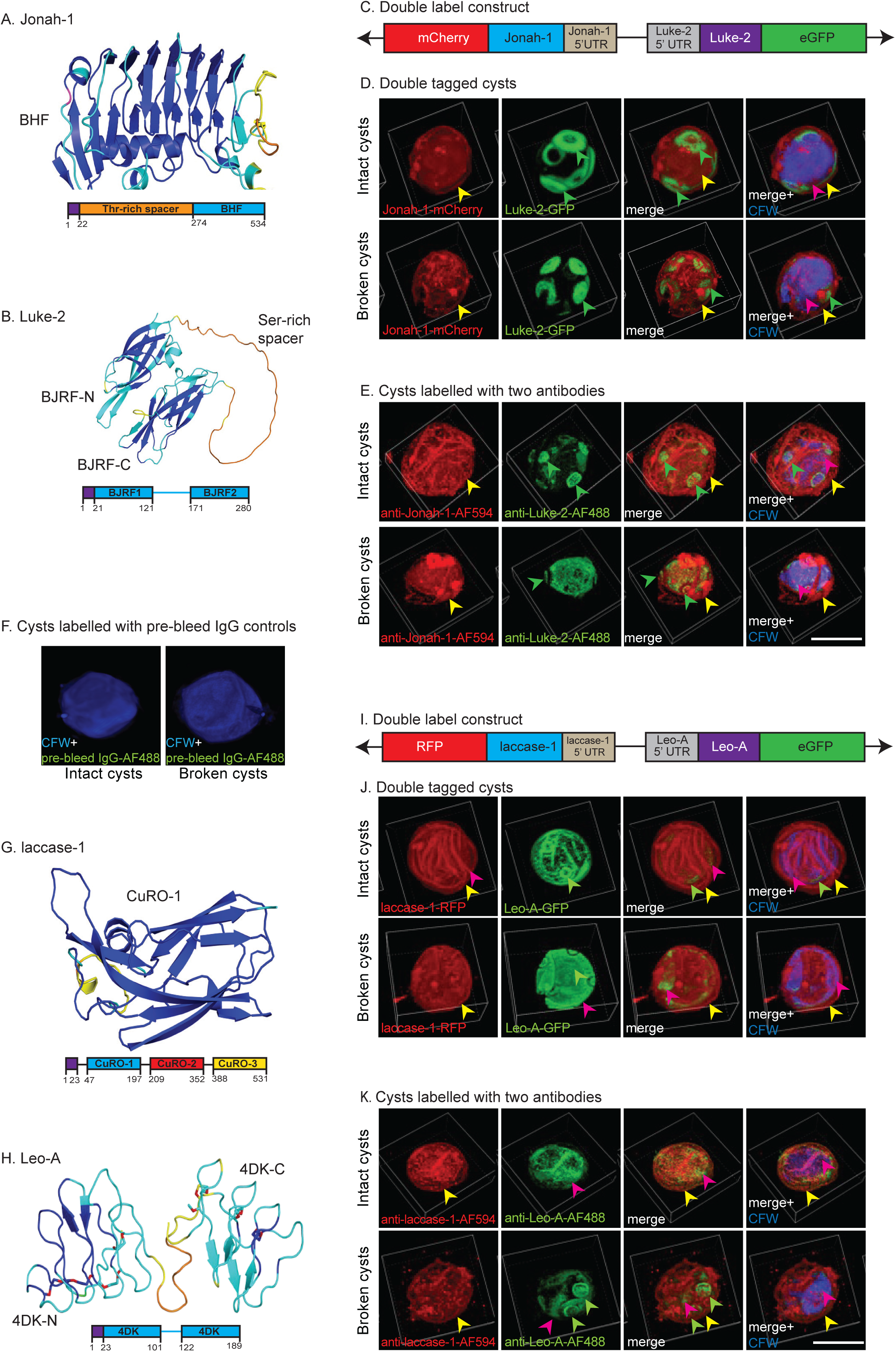
Double labels show rabbit antibodies (rAbs) to Jona-1, Luke-2, laccase-1 and Leo-A localize to similar places in intact cysts and broken cyst walls as do tagged proteins expressed under their own promoter in transfected Ac. A. AlphaFold with confidence colored shows the single BHF, which was used to make the EcMBP-Jonah-1 fusion-protein for production of anti-Jonah-1 rAbs. The diagram shows the N-terminal signal peptide (purple) and low complexity Thr-rich domain (orange). B. Alphafold shows N- and C-terminal BJRFs (BJRF-N and BJRF-C) connected by a low complexity spacer, which was used to make the EcMBP-Luke-2 fusion-protein to produce anti-Luke-2 rAbs. The diagram shows the N-terminal signal peptide (purple) and the low-complexity Ser-rich domain. C. The first plasmid for double-labeling cyst walls contains full-length Jonah-1 cDNA with 574-bp of the 5’ UTR and a C-terminal mCherry tag head-to-head with full-length Luke-2 cDNA with 446-bp of 5’ UTR and a C-terminal GFP tag (38). D. Double labels show Jonah-1-mCherry is present in the ectocyst layer (yellow arrowheads) of intact and broken cysts, while Luke-GFP predominates in the ostioles (green arrowheads) of both preparations. CFW marks the endocyst layer (red arrowheads). E. Anti-Jonah-1 rAbs (red) bind to the ectocyst layer of intact and broken cysts, while anti-Luke-2 rAbs (green) bind to ostioles of intact cysts and to the endocyst layer of broken walls. F. Rabbit pre-bleeds fail to bind to intact cysts and broken walls. G. AlphaFold shows the first copper oxidase domain (CuRO-1), which was used to make EcMBP-CuRO-1 for production of anti-laccase-1 rAbs. The diagram shows three CuRO domains in laccase-1. H. Alphafold shows N- and C-terminal 4DKs (4-DK-N and 4DK-C), which were used to make EcMBP-Leo-A for production of anti-Leo-A rAbs. The diagram shows the short spacer between 4DKs of Leo-A. I. The second plasmid for double-labeling cyst walls contains full length laccase-1 cDNA with 405-bp of the 5’ UTR and a C-terminal RFP tag head-to-head with full length Leo-A cDNA with 486-bp of the 5’ UTR and a C-terminal GFP. J. Laccase-1-RFP is in the ectocyst layer of intact and broken cysts, while Leo-A-GFP is present in the endocyst layer and ostioles. K. Anti-laccase-1 rAbs (red) bind to the ectocyst layer of intact and broken cysts, while anti-Leo-A rAbs (green) bind to the endocyst layer of intact cysts and to ostioles in broken walls. Scale bars for D-F, J, and K are 5 µm.

The caveat here is the CuRO-1 domain of Ac laccase-1, which has sequences (TMWYHDH and NVYAGLAGFYLLRD) conserved in two other Ac laccases (ACA1_006180 and ACA1_008840) and partially conserved in laccases of bacteria (e.g. *Clostridium* and *Bacillus*), fungi (*Bifiguratus*), and plants (*Daucus*) (Fig. S1) (56, 68, 69). These results, which suggest the possibility that anti-laccase-1 rAbs might cross-react with bacteria, fungi, or plant cells in the eye, are not surprising, because laccase is a well-conserved enzyme (70). TBLASTN showed that all *Acanthamoeba* genomes in AmoebaDB have >90% positional identities with Curo-1 of Neff strain and so are likely to efficiently bind anti-laccase-1 rAbs.

### Confocal microscopy of rabbit antibodies (rAbs) binding to intact cysts and broken walls of the Ac Neff strain generally confirmed the localization of tagged wall proteins

The goal here was to determine whether rAbs to four wall proteins localize to the same place in cyst walls of the Neff strain of Ac as do tagged proteins expressed under their own promoters (38, 40). We used confocal microscopy to compare the binding of pairs of rAbs to broken and intact walls of non-transfected Neff strain cysts with the localization of pairs of tagged proteins, each expressed under its own promoter, in walls of transfected Ac (Figs. 1C and 1I). Double labels showed Jonah-1-mCherry has a patchy distribution in the ectocyst layer of intact and broken cyst walls of Ac, while Luke-2-GFP is more abundant in ostioles than the endocyst layer, which was labeled with CFW (Fig. 1D). Similarly, rAbs to the BHF of Jonah-1 labeled red the ectocyst layer of intact and broken cyst walls, while rAbs to two BJRFs of Luke-2 labeled green the ostioles of intact cysts and the endocyst layer of broken cysts (Fig. 1E). In contrast, the negative control, which is pre-bleed rAbs, did not bind to intact or broken cysts (Fig. 1F).

Laccase-1-RFP had a patchy distribution in the ectocyst layer of intact and broken cyst walls of Ac, while Leo-A-GFP was more abundant in the endocyst layer than the ostioles (Fig. 1J). The rAbs to Curo-1 domain of laccase-1 labeled the ectocyst layer of both intact cyst and broken walls, while rAbs to two 4DKs of Leo-A more heavily labeled the endocyst layer of intact cysts and the ostioles of broken walls (Fig. 1K). In summary, localizations of rAbs to wall proteins matched well localization of tagged proteins.

### Despite numerous unexpected binding patterns, rAbs to Jonah-1, Luke-2, laccase-1, and Leo-A all efficiently detected CFW-labeled cysts of 10 of 11 *Acanthamoeba* isolates obtained from the ATCC

The goals here were to determine whether rAbs to cyst wall proteins bind to the same place in cyst walls of 10 other *Acanthamoeba* isolates as in the wall of the model Neff strain of Ac and to quantitatively measure how efficiently the rAbs detect CFW-labeled cysts. Eight *Acanthamoeba* isolates from the ATCC including T4, T7, T11, and T18 genotypes, were a generous gift from Dr. Monica Crary of Alcon, Inc., who studies their binding to contact lenses (44, 47). A T4 genotype and another AK isolate (Esbc4) came from Noorjahan Panjwani, who identified AcMBP-1 on the surface of trophozoites (24–27). Segments of 18S RNA genes were amplified from trophozoites, cloned, and sequenced to confirm their identity prior to encystation and testing with rAbs (Fig. S2) (16–21, 67).

High-power confocal microscopy showed rAbs to the BHF of Jonah-1 bound in a somewhat patchy distribution to the ectocyst layer of cysts of Neff strain and 10 other *Acanthamoeba* isolates from the ATCC (Figs. 1A and 2A). In contrast, rAbs to CuRO-1 domain of laccase-1 bound in a homogenous pattern to the Neff strain and to other *Acanthamoeba* isolates, which is like that of laccase-1-GFP (Figs. 1G, 1J and 2B). While rAbs to Luke-2 and Leo-A densely labeled the ostioles and weakly labeled the endocyst layer of cysts of the Neff strain and a few ATCC isolates (Figs. 1B, 1H, 2C and 2D), most other cysts were labeled on the ectocyst layer. There are two explanations, which are not mutually exclusive, for the variable distribution in the cyst walls of rAbs to Luke-2 and Leo-A. First, these proteins may be made at different times by different *Acanthamoeba* isolates, so that Luke-2 and Leo-A localize to different places (38). Second, these rAbs may bind to relatively small amounts of Luke-2 and Leo-A in the ectocyst layer, which were not visualized by confocal microscopy, because of the large amounts of GFP-tagged Luke-2 and Leo-A in the endocyst layer and ostioles (Figs. 1D and 1J).

**Fig. 2.**
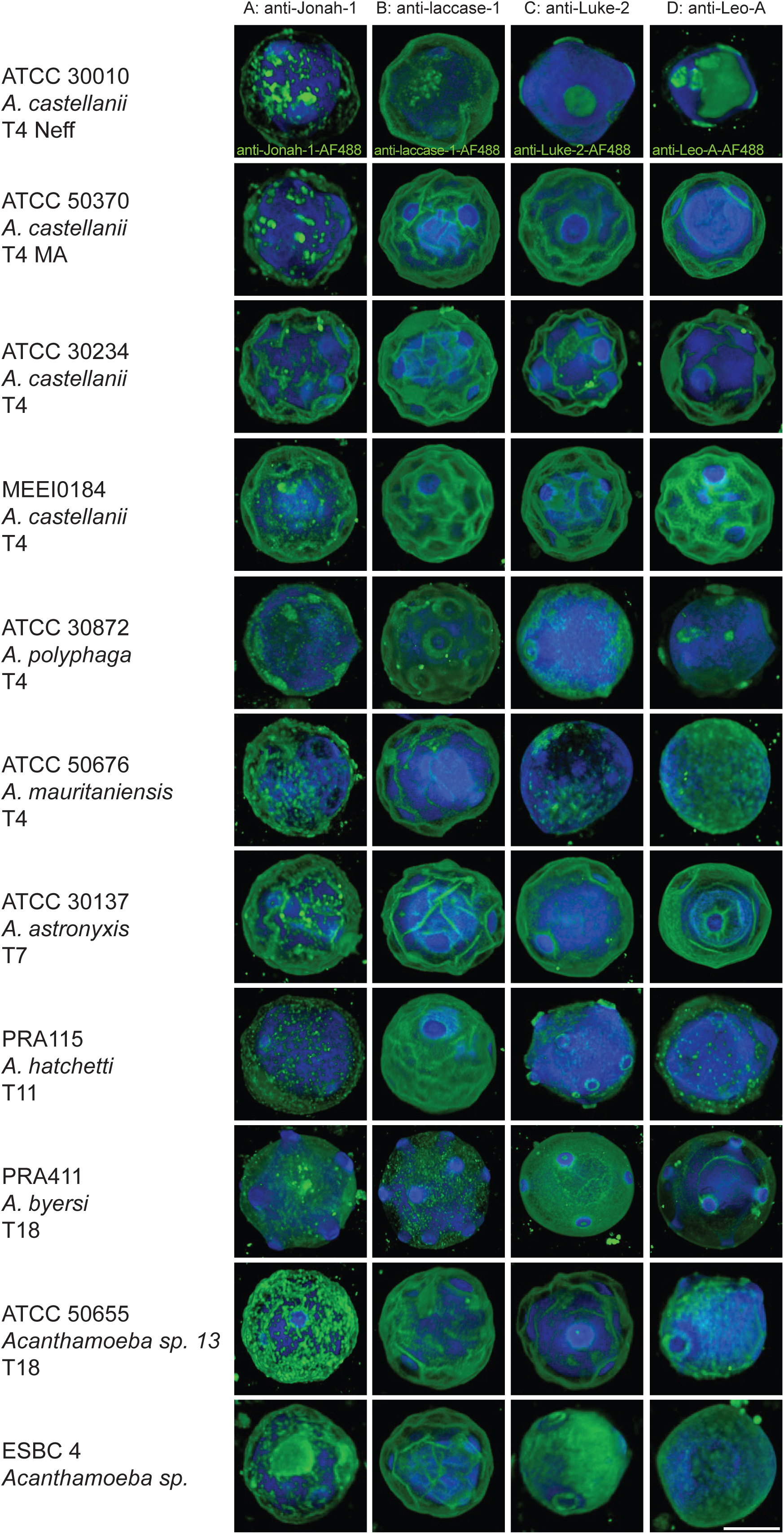
High power confocal microscopy of 11 *Acanthamoeba* isolates from the ATCC shows anti-Jonah-1 and anti-laccase-1 rAbs consistently bind to the surface of cysts, while anti-Luke-2 and anti-Leo-A rAbs bind to the endocyst layer and ostioles of some cysts (expected) and to the ectocyst layer of other cysts (unexpected). A and B. The anti-Jonah-1 rAbs bind in a patchy distribution to the surface of cysts of 11 *Acanthamoeba* isolates from the ATCC, while the anti-laccase-1 rAbs bind in a homogenous manner. CFW labels the endocyst layer. C and D. Both the anti-Luke-2 and anti-Leo-A rAbs bind to the ostioles and endocyst layer of some cysts (e.g., *A. polyphaga, A. hatchetii, A, byersi*, and ESBC4) and the ectocyst layer of other cysts (e.g., three *A. castellanii* and *A. astronyxis*). Possible reasons for the different patterns of binding of rAbs to Luke-2 and Leo-A are discussed in the Results. A single scale bar for A to D is 5 µm. Low power views of rAbs binding to 11 *Acanthamoeba* species are shown in Fig. S3, while percentages of CFW-tagged cysts detected are shown in Fig. 3A.

To determine which wall proteins might be the best targets for diagnosis of cysts in AK, we counted in randomly selected low power confocal fields the percentage of CFW-labeled cysts detected by rAbs to the four wall proteins (Figs. 3A and S3). The rAbs to Luke-2, Leo-A, and laccase-1 each detected >95% of CFW-labeled cysts of 10 of 11 *Acanthamoeba* isolates tested. In contrast, anti-Jonah-1 rAbs showed >95% detection for just 7 of 11 *Acanthamoeba* isolates, suggesting that Jonah-1 might not be quite as good a target. In addition, all four rAbs struggled to detect cysts of *A. mauritaniensis*, suggesting that there was a problem with encysting these trophozoites under the conditions used here. While counts were performed for two sets of cysts labeled with Protein-A Sepharose-purified antibodies, we obtained similar results with rabbit sera diluted 1:300. Most important, we confirmed that rAbs to all four proteins were also visible with a conventional fluorescence microscope, which is like those present in clinical labs or in ophthalmologists’ offices (Fig. S4).

**Fig 3.**
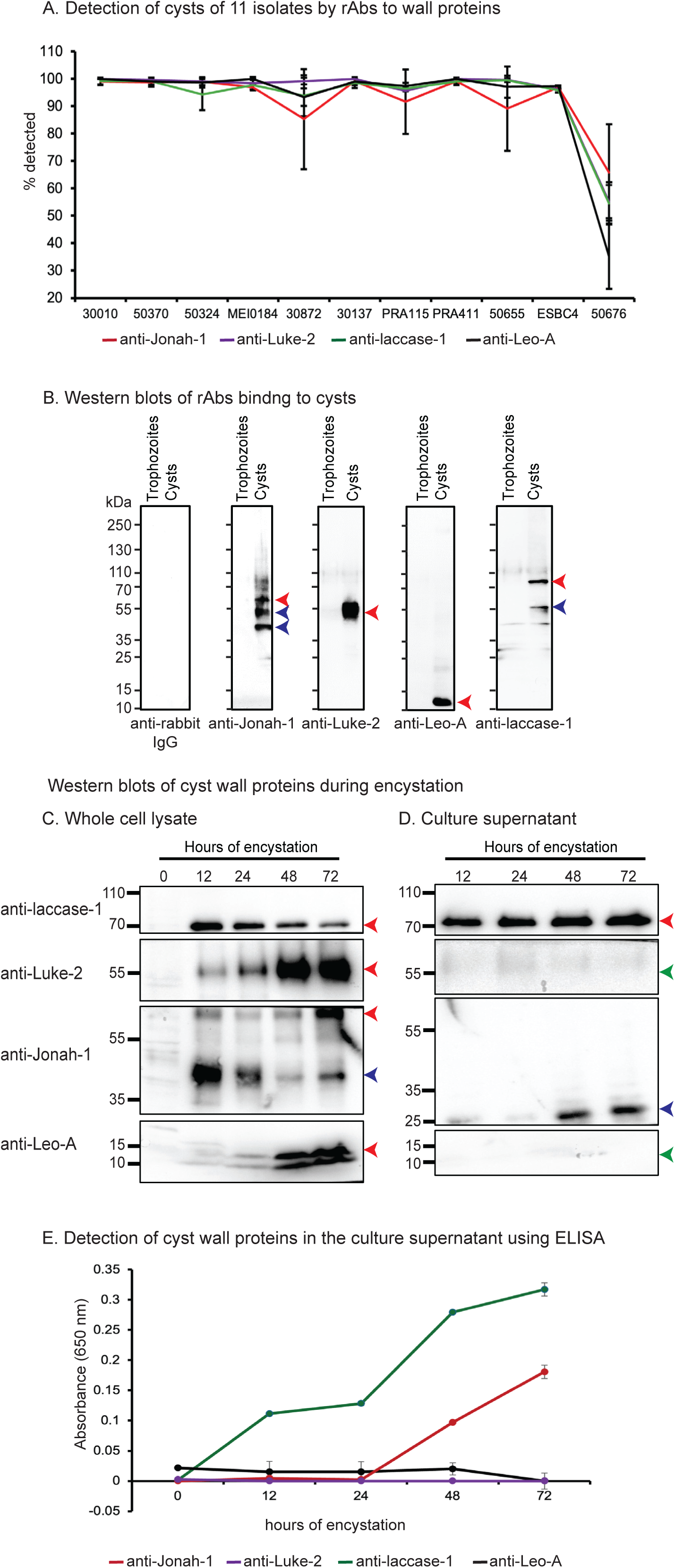
Plots show rAbs to Jonah-1, laccase-1, Luke-2, and Leo-A all efficiently detect CFW-labeled cysts, while Western blots and ELISAs identify laccase-1 shed into the medium of encysting Ac. A. The plot shows the percentage of CFW-labeled cysts detected by four rAbs in two independent experiments (average plus SEM), in which 100+ cysts were counted in random low power fields (Fig. S3). The rAbs to Luke-2, Leo-A, and laccase-1 each detect >95% of cysts of 10 of 11 *Acanthamoeba* isolates tested, while the rAbs to Jonah-1 are slightly less efficient in detecting three isolates of *Acanthamoeba*. The exception is *A. mauritaniensis* (50676), which was poorly detected with all four rAbs, suggesting a problem with cyst formation. B. Western blots show pre-bleeds from rabbits (negative control) fail to bind to proteins of trophozoites or cysts, while rAbs to Jonah-1, Luke-2, Leo-A, and laccase-1 fail to bind to trophozoite proteins (a second negative control) but bind well to cyst proteins of the Neff strain. As discussed in detail in the Results, the anti-Jonah-1 rAbs bind to the expected 55-kDa band (red arrowhead) and to two lower mol wt bands (blue arrowheads), while the anti-Luke-2 rAbs bind to a heavy and broad 50-kDa band (red arrowhead), which is greater than the expected size of 27-kDa. The anti-Leo-A rAbs bind to a 13-kDa band (red arrowhead), which is slightly smaller than the expected size of 17-kDA, while the anti-laccase-1 rAbs bind to a 75-kDa band (red arrowhead), which is slightly bigger than the expected size of 64-kDa, as well as to a less abundant 55-kDa band (blue arrowhead). C. Western blots of whole cell lysates with rAbs show laccase-1 and Jonah-1 are made early in encystation, while Luke-1 and Leo-A are made later in encystation. These results are consistent with the expression of tagged proteins (38). D. Western blots of culture supernatants show that intact laccase-1 is abundant throughout encystation, while a degraded form of Jonah-1 is abundant at 48- and 72-hour encystation. In contrast, neither Luke-2 nor Leo-A are present in the culture medium (green arrowheads). E. Consistent with Western blots, direct ELISAs of culture supernatants detected laccase-1 throughout encystation; Jonah-1 was detected at 48- and 72-hours encystation, while neither Luke-2 nor Leo-A were detected.

### Western blots showed rAbs to recombinant Jonah-1, Luke-2, Leo-A, and laccase-1 bind to native wall proteins of roughly the expected sizes and timing of expression, while enzyme-linked immunoassays (ELISAs) detected intact laccase-1 shed into the medium by encysting *Acanthamoebae*

In addition to confirming the size and timing of expression of each wall protein recognized by its rAbs, the goal here was to determine if any of the wall proteins are secreted into the culture medium during encystation and so might be detected using an ELISA. Western blots showed that pre-bleeds from rabbits do not react with proteins of trophozoites or cysts of the Neff strain of Ac (Fig. 3B). While rAbs to MBP-fusions with Jonah-1, Luke-2, Leo-A and laccase-1 did not bind to trophozoite proteins, all rAbs bound well to cyst proteins (Fig. 3B). Anti-Jonah-1 rAbs bound to a protein of the expected size of ∼55-kDa (red arrow), as well as to two lower molecular weight bands (blue arrows), which may result from proteolytic cleavage of Jonah-1. Anti-Luke-2 rAbs bound to a thick ∼50-kDa band (red arrow), which is greater than the 27-kDa expected size of Luke-2. The increased size of Luke-2 is likely caused by extensive glycosylation of its five *N*-glycan sites and ∼20 *O*-glycan sites in the low complexity, Ser-rich spacer. Anti-Leo-A rAbs bound to an ∼13-kDa band (red arrow), which is slightly smaller than the expected size of 17-kDa, most likely due to the acidity of the protein that makes it run faster on SDS-PAGE. Anti-laccase-1 rAbs bound to a 75-kDa band (red arrow), which is slightly bigger than the expected size of 64-kDa and so likely caused by glycans, as well as to a less abundant 55-kDa band (blue arrow) likely caused by proteolytic cleavage.

Western blots of proteins of encysting Ac showed Jonah-1 and laccase-1 are each made early, while Luke-2 and Leo-A are each made later, which is consistent with observations of GFP-tagged proteins (Fig. 3C) (38, 40). Western blots of proteins secreted into the culture medium by encysting Ac showed intact laccase-1 and a degraded form of Jonah-1, while Luke-2 and Leo-A were absent (Fig. 3D). Laccase-1 was detected by a direct ELISA in the culture medium beginning at 12 hours encystation; Jonah-1 appeared later and was less abundant, while Luke-2 and Leo-A were not detected (Fig. 3E).

### Structural and experimental evidence that an antiparallel β-sandwich (ABS) is the mannose-binding domain (ManBD) of the *Acanthamoeba castellanii* mannose-binding protein (AcMBP-1)

While the primary goal here was to identify a simple target for diagnosis of *Acanthamoeba* trophozoites in cultures of corneal scrapings, the secondary goal, which is presented first, was to determine the structure and experimentally test the ManBD of AcMBP-1 that mediates adherence to corneal epithelial cells (24–27). AcMBP-1, which is 833-aa long, was identified in the MEEI 0184 isolate of *Acanthamoeba* (AAT37864 in the NR database at NCBI) but is absent from AmoebaDB (although we identified it in our unpublished transcriptome and proteome of Neff strain of Ac) (16, 17). A recent sequence analysis of AcMBP-1 showed it contains an N-terminal DUF4114 domain, followed by a large Cys-rich region (CRR), a transmembrane helix, and a ∼70-aa cytosolic domain (23, 69). AcMBP-2, which is 360-aa long and is present in AmoebaDB (ACA1_248600), contains the DUF4114 domain but is missing most of the CRR. Structure predictions of AcMBP-1 showed that the DUF4114 domain is an antiparallel β-sandwich (ABS), while the CRR is composed of many pairs of antiparallel β-strands secured at both ends by disulfides (Fig. 4A) (60, 71). Structure searches showed the ABS of AcMBP-1 closely matches that of an uncharacterized protein of *Myxococcus sp.* (30% identity over a 207 amino acid overlap with an e-value of 1.4e-13) (Fig. 4B), as well as the ABS of the BclA lectin of *Burkholderia cenocepacia* that has been crystallized with bound methyl α-D-mannoside (12% identity over a 157 amino acid overlap with an e-value of 2.3e-2) (Fig. 4C) (63, 72). While the ABSs of the bacterial proteins lack Cys residues, the ABS of AcMBP-1 has four sets of disulfide bonds, which link short loops that are reminiscent of 4DKs of Leo lectins (Fig. S5) (38).

**Fig. 4.**
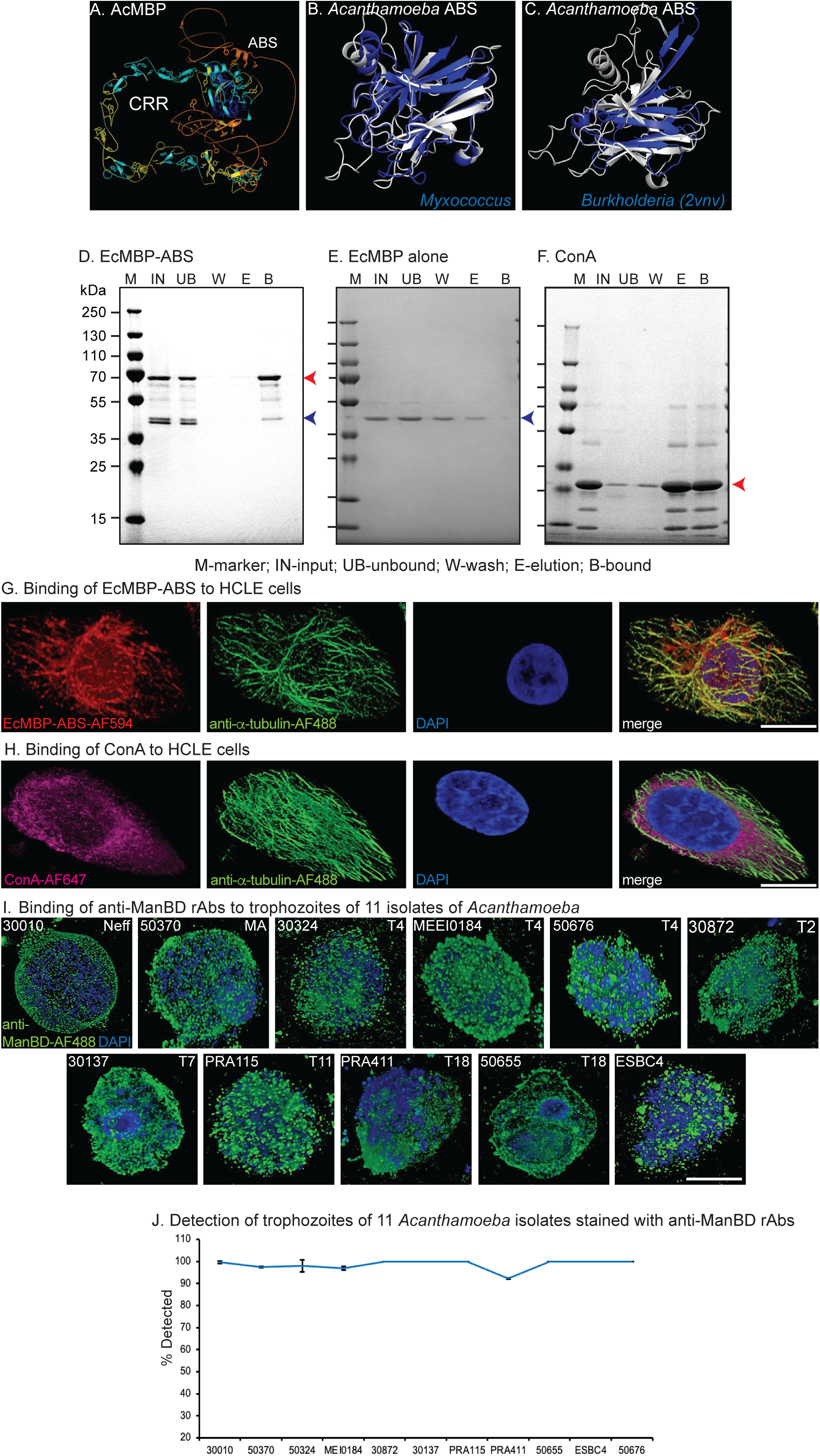
Experimental evidence that the antiparallel β-sandwich (ABS) of AcMBP is the ManBD and demonstration that anti-ManBD rAbs efficiently detect DAPI-labeled trophozoites of all 11 *Acanthamoeba* species. A. AlphaFold with the confidence-colored shows AcMBP contains an N-terminal ABS and a large CRR composed of anti-parallel β-strands linked by disulfides at each end. B. Foldseek and PyMOL show that the ABS of AcMBP closely matches the same domain in an uncharacterized protein of *Myxococcus*. C. The ABS also resembles the mannose-binding domain of *Burholderia*, which has been crystallized (PDB 2vnv), although the match is less strong. D. A Western blot shows an EcMBP-ABS fusion-protein binds so strongly to a mannose-agarose resin that it cannot be eluted with excess α-methyl-mannose but can only be released with SDS. The negative control EcMBP alone (E) fails to bind to the mannose-agarose resin, while the positive control ConA (F) is eluted from the mannose-agarose resin with excess α-methyl-mannose. Confocal microscopy shows EcMBP-ABS (G) and ConA (H) each bind to an HCLE cell labeled with an anti-tubulin antibody and DAPI. See also Fig. S6 for low-power views of EcMBP-ABS and ConA binding to HCLE cells, as well as the failure of EcMBP alone (negative control) to bind to HCLE cells. I. The rAbs to EcMBP-ABS, renamed here ManBD, binds to the surface of DAPI-labeled trophozoites of 11 *Acanthamoeba* species, which were used for binding anti-cyst antibodies in Fig. 2. J. Counts of low-power views of trophozoites (Fig. S6) show that >90% of DAPI-labeled trophozoites were detected by rAbs to the ManBD. Scale bars for G to I are each 5 µm.

Experimental evidence that the ABS is the ManBD of AcMBP-1 included binding of most of an EcMBP-ABS fusion-protein to a mannose-agarose resin, while there was minimal binding of EcMBP alone (blue arrow) (negative control) (Figs. 4D and 4E). Of note here are 1) the full-length EcMBP-ABS fusion-protein (red arrow) bound best to mannose-agarose resin, while a degradation product (blue arrow) bound very weakly and 2) EcMBP-ABS failed to elute with excess α-methyl-mannose, suggesting binding is very tight. In contrast, the mannose-binding plant lectin Concanavalin A (ConA) (red arrow) (positive control) bound well to the mannose-agarose resin and was eluted with α-methyl-mannose, suggesting binding is less tight (Fig. 4F). EcMBP-ABS bound to the surface and to vesicles associated with microtubules of human corneal limbal epithelial (HCLE) cells (Figs. 4G and S6). ConA bound in a similar pattern to HCLE cells, and there was no binding of EcMBP alone (Figs. 4H and S6). In summary, structural and experimental evidence suggests that ABS domain of AcMBP-1 binds mannose, and so we now refer to it as the ManBD.

### Anti-ManBD rAbs efficiently detected trophozoites of all 11 *Acanthamoeba* isolates from the ATCC

BLASTP showed there are many identical amino acid sequences in the ManBDs of AcMBP-1 and AcMBP-2, so that it is likely and our anti-ManBD rAbs will cross react with both proteins. Similarly, TBLASTN of *Acanthamoeba* genomes in AmoebaDB showed the vast majority have >85% positional identity with ManBD, strongly suggesting that anti-ManBD rAbs will bind well to trophozoites of all *Acanthamoeba* species. Indeed, anti-ManBD rAbs, which were made by immunizing a rabbit with the EcMBP-ABS fusion-protein, bound to the surface of trophozoites of the Neff strain, while pre-bleed rabbit sera failed to bind to trophozoites (Figs. 4I and S6). High power confocal micrographs showed the anti-ManBD rAbs bound to trophozoites of all 11 *Acanthamoeba* isolates from the ATCC (Fig. 4I). DAPI-labeled trophozoites appear blue because *Acanthamoeba* mitochondria have a relatively large genome (∼50-kDa each) (73). Counts of low power confocal micrographs showed that anti-ManBD rAbs efficiently detected DAPI-labeled trophozoites (Figs. 4J and S6). Finally, we confirmed that anti-ManBD rAbs were also visible with a conventional fluorescence microscope like those used in clinical settings (Fig. S4).

## DISCUSSION

### Major conclusions and their limitations

Structural predictions and searches not only helped us identify and characterize cellulose-binding domains of Luke-2 and Leo-A (38) and the ManBD of AcMBP-1 (performed here) but made it easy to prepare large quantities of recombinant proteins used to immunize rabbits for polyclonal antibodies. Remarkably, abundant wall proteins do not have to be in the ectocyst layer (Jonah-1 and laccase-1) but may be predominantly in the endocyst layer (Luke-2 and Leo-A) and still be excellent diagnostic targets for cysts. Further, the binding of rAbs to Jonah-1, Luke-2, Leo-A, and laccase-1 is great enough, so that cysts were easily detected with a conventional fluorescence microscope used in offices of eye doctors or clinical labs. Finally, laccase-1 is shed by encysting *Acanthamoebae*, suggesting the possibility that a lateral flow immunoassay might be used for point-of-care testing (74). Diagnostic anti-cyst rAbs will complement monoclonal antibodies to AcMBP-1 (24, 26–28), as well as antibodies to transporters and secreted proteins (75–77). Anti-cyst antibodies may also supplement Loop-mediated Isothermal Amplification (LAMP) assays for diagnosing AK (78–80).

The most important limitation here is that the anti-cyst rAbs have not yet been tested using confocal microscopy of eyes or corneal scrapings from patients suspected of having AK. Although we are confident that Luke-2, Leo-A, and Jonah-1 antigens are each unique to Ac, anti-laccase-1 rAbs may cross-react with walls of bacteria, fungi, or plants. Cysts examined here were made by starving cultured *Acanthamoebae* and so might not be the same as those made in the corneal epithelium, soil, or water. Further, for commercialization, rAbs will have to be replaced by mouse monoclonal antibodies, which are used for stool diagnosis of walled forms of *Giardia*, *Entamoeba,* and *Cryptosporidium*, or by nanobodies that are used for diagnosis of viruses and bacteria (81–84).

While the DUF4114 had been suggested as the ManBD of AcMBP-1 (23), we used structural predictions, recombinant protein expression, and binding to mannose-agarose resins and HCLE cells to demonstrate that the ABS of AcMBP-1 is indeed the ManBD. Further, we showed that anti-ManBD rAbs to efficiently identify trophozoites from all 11 ATCC isolates of *Acanthamoebae*. A limitation of our characterization of the ManBD of AcMBP-1 is that we have not performed Ala mutations to identify the amino acids involved in binding mannose on host cell glycoproteins, nor has the ManBD been tested versus arrays of mammalian carbohydrates to determine the precise linkages of terminal mannose to *N*- or *O*-linked glycans (85). Again, rAbs to ManBD need to be tested versus trophozoites cultured from eye scrapings of patients suspected of having AK, while monoclonal antibodies or nanobodies need to be made for the commercialization of anti-ManBD diagnostics.

## MATERIALS AND METHODS

### Ethics statement

Culture and manipulation of *Acanthamoebae* under BSL-2 protocols were approved by the Boston University Institutional Biosafety Committee. Production of custom rabbit antibodies was approved by the Institutional Animal Care and Use Committee of Cocalico Biologics, Inc., Denver PA.

### Summary of new methods to study candidate targets for diagnosis of *Acanthamoeba* cysts and trophozoites

Methods for culture and encystation of *Acanthamoebae*, protein structure predictions, expression of pairs of tagged cyst wall proteins in transfected parasites, production of EcMBP-fusions in the periplasm of *E. coli*, and confocal microscopy were all described in detail in our recent study of the roles of cellulose-binding and timing of expression in targeting of proteins to the cyst wall of the Neff strain of Ac (38). Here structure predictions were used to identify a candidate ManBD in the AcMBP, which was tested by binding of an EcMBP-fusion to a mannose-agarose resin and to HCLE cells. We made rAbs to EcMBP-fusions of four wall proteins and the ManBD and determined their binding to cysts and to trophozoites, respectively. The efficiency of detecting CFW-labeled cysts and DAPI-labeled trophozoites was quantified for 11 *Acanthamoeba* isolates from the ATCC, the identities of which were confirmed by sequencing 18S RNAs. Finally, unless stated otherwise, each experiment was repeated at least three times.

### *Acanthamoeba* genotypes, culture, and cyst preparation

Trophozoites of the Neff strain of Ac (ATCC 30010) were obtained from the American Type Culture Collection (ATCC). Trophozoites of other strains of *Acanthamoeba*, originally derived from human corneal infections and granulomatous encephalitis infections, were acquired from Dr. Monica Crary, Alcon Research, LLC, Fort Worth, TX, United States, or from Noorjahan Panjwani of Tufts University Medical School (26, 44). Each of these *Acanthamoeba* isolates was confirmed by PCR of 18S RNA, the sequences of which are shown in Fig. S3. Trophozoites were grown and maintained in axenic culture at 30°C in T-75 tissue culture flasks in 10 ml ATCC medium 712 (PYG plus additives) with antibiotics (Pen-Strep) (Sigma-Aldrich Corporation, St. Louis, MO), as described previously (38, 40, 86). Cysts were prepared from trophozoites by incubating them with an encystation medium (EM, 20 mM Tris-HCl [pH 8.8], 100 mM KCl, 8 mM MgSO_4_, 0.4 mM CaCl_2,_ and 1 mM NaHCO_3_) for 120 hours(87).

### Production of EcMBP-fusions containing four cyst wall proteins and the ABS of AcMBP

The production and characterization of EcMBP-fusions containing the BHF of Jonah-1, two BJRFs of Luke-2, two sets of 4DKs of Leo-A, and the CuRO-1 of laccase-1 have all been described (38). The structure of AcMBP was predicted by AlphaFold2 using the Colab server (60), while the molecular visualization of the ABS of AcMBP, including its four disulfide knots, was performed using PyMOL Molecular Graphics System, Version 2.0 Schrödinger, LLC (88). The ABS structure was then compared with AlphaFold and PDB databases using the Foldseek server (63). Finally, the ABS was made into an EcMBP fusion protein, using the same methods to make EcMBP-fusions containing wall proteins. EcMBP-fusions were made in the pMAL-p2x vector (New England Biolabs) and expressed in BL21-CodonPlus (DE3)-RIPL cells (Agilent Technologies, Lexington, MA) (65, 66). The overexpression of MBP-fusion proteins was induced by the addition of 0.1 mM of IPTG for 16 hours at 16°C. MBP-fusion proteins were affinity purified with amylose resin following the manufacturer’s instructions (New England Biolabs). The identity and purity of recombinant purified MBP-fusion proteins were confirmed by SDS-PAGE and western immunoblotting analyses.

### Generation, IgG purification, and Alexa Fluor labeling of rabbit antibodies (rAbs)

Purified EcMBP-fusions containing four wall proteins and the ABS of AcMBP were used to raise custom rabbit polyclonal antibodies, as per the standard protocol of Cocalico Biologicals, Denver, PA. Total IgG was purified from plasma samples of pre-immune and post-immunized rabbits using affinity chromatography (Pierce™ Protein A Agarose, Thermo Fisher Scientific, USA). In brief, the serum was first diluted two-fold with binding buffer (1x Tris-Buffered Saline, pH 7.4) and loaded onto the top of a column containing Protein A Agarose beads, which were washed with 1x TBS (20-fold column volume). The bound IgG was eluted with 0.1 M glycine-HCl (pH 2.7) into the neutralizing buffer (1 M Tris-HCl, pH 9.0) aliquoted in advance. The elution was concentrated, and the buffer was exchanged against 25 mM Tris-HCl 150 mM NaCl (pH 8.0) using a 30 kDa Amicon Ultra centrifugal filter (Millipore, USA). The purified rAbs were either used as such or further labeled with Alexa flour dyes (i.e. 488 and 647) using Invitrogen^Tm^ Fluorescent Protein labeling Kits (Thermo Fisher Scientific), as per the manufacturer’s instructions.

### Confocal microscopy to compare the localizations of tagged proteins in cyst walls to those of bound rAbs to wall proteins and to determine the efficiency of detecting CFW-tagged cysts with rAbs

The localization of pairs of wall proteins (Jonah-1-mCherrry and Luke-2-GFP or laccase-1-RFP and Leo-A-GFP), each expressed under its own promoter, were previously shown by confocal microscopy of transfected Neff strain cysts (38). Here the localizations of tagged proteins in intact cysts and walls broken by sonication were compared with localizations of pairs of rAbs (anti-Jonah-1-AF594 and anti-Luke-2-AF488 or anti-laccase-1-AF594 and anti-Leo-A-AF488) binding to intact cysts and broken walls. Intact cysts and broken walls were collected by centrifugation, washed in PBS, fixed in 4% paraformaldehyde for 15 minutes at room temperature, washed again with PBS, and blocked with 1% BSA for one hour at room temperature. Cysts and walls were then incubated with pairs of tagged rAbs (1:200 dilution) conjugated with AF488 or AF594 for one hour at room temperature, washed with PBS, and incubated with CFW 1:20 (CFW, 1 mg/ml, Sigma Aldrich) for 30 minutes at room temperature. The cells were washed three times with 1x PBS and mounted in VECTASHIELD® Antifade Mounting Medium (Vector Laboratories, Newark, CA). Cysts were imaged using CFI Plan Apochromat VC 60XC NA 1.42 oil objective of Nikon Ni2 AX inverted confocal microscope equipped with FX-Format F-mount cameras Digital Sight 10 and Digital Sight 50M. We deconvolved 0.1 μm optical sections using NIS elements (Version: AR5.41.02) imaging software. All confocal images shown were 3D reconstructions using dozens of z-stacks. Size bars were based upon 2D cross-sections.

To determine how well rAbs to four wall proteins detect cysts of 11 isolates of *Acanthamoeba* from the ATCC, mature cysts (120 hours in encystation medium) from each isolate were collected, washed in PBS, fixed in 4% washed again, and blocked with BSA, as described above. The cysts were incubated with unlabeled rAbs to each wall protein, which was detected as secondary anti-rabbit IgG conjugated with Alexa flour 488 (1:300) and CFW. Confocal images were captured with the 60X objective and 3D reconstructions were performed to determine whether rAbs bound to the ectocyst layer, endocyst layer, and/or ostioles. In addition, at least 100 cysts in random fields were counted to determine what percentage of CFW-labeled cysts of each isolate were detected with each rAb. Each experiment was repeated at least twice and counts for the two experiments were averaged. Finally, we examined the same set of slides with a Plan-APO Chromat 100x oil immersion lens on a Zeiss Axio Observer Z1 microscope equipped with Axio Cam ERc5s to show that a confocal microscope was not needed to detect rAbs-labeled trophozoites or cysts.

### Western blotting and ELISA of wall proteins in trophozoites, cysts, and medium of encysting *Acanthamoebae*

To detect the cyst wall proteins in the Neff strain of Ac, log-phase trophozoites and mature cysts (120 hours in encystation medium) were harvested and lysed in an SDS sample buffer. The lysates were separated on SDS-PAGE gel (4-15%), transferred to a nitrocellulose membrane, and blocked with 5% BSA in PBS. The blots were probed with rAbs (1:5000) or purified rabbit IgG (1:1000) raised against the different abundant cyst wall proteins. Anti-rabbit IgG conjugated to HRP (Thermo Fisher Scientific) was used as the secondary antibody. Rabbit pre-immune serum or anti-rabbit IgG was used as a control. Super Signal West Pico PLUS (Thermo Fisher Scientific) substrate was used for chemiluminescent detection. Blots were imaged using GE ImageQuant LAS 4000 gel imager.

To detect cyst wall proteins in encysting Ac and their culture supernatant, trophozoites of the Neff strain were encysted and the cell pellet and culture supernatant were collected at 0, 12, 24, 48, and 72 hours. To remove the possibility of intact cells, the supernatant was precleared by three rounds of centrifugation at 3000xg for 10 min at 4℃. The supernatant was subjected to an additional centrifugation step (10000g, 10 min at 4℃) and concentrated using a 3 kDa Amicon Ultra centrifugal filter (Millipore, USA). The protein concentration in each sample was determined by DC protein assay (Bio-Rad, Hercules CA, 500–011). The presence of candidate cyst wall proteins in concentrated culture supernatant was studied by western blotting and direct ELISA methods. For western blotting analysis, a total of 50 µg protein from each time points were separated on SDS-PAGE gel (4-20%), transferred to nitrocellulose membrane, and probes, as described above. For direct ELISA, the flat bottom microtiter plates (Nunc-Immuno™ MicroWell™ 96 well solid plates) were coated overnight with 50 µl of the culture supernatants diluted to a final concentration of 50 µg/ml in carbonate buffer. Negative controls were carbonate buffer alone or trophozoite culture medium. The microtiter plates were blocked with 200 µl blocking buffer (3% BSA/PBS) at room temperature for two hours and washed three times with PBS/Tween-20. The plate was subsequently incubated with either control pre-immune rabbit IgG or immunized rabbit purified IgG for two hours, washed, and incubated with anti-rabbit HRP conjugated secondary antibody. Plates were washed and then developed using TMB ELISA substrate (Abcam, Ab171522), and the absorbance was recorded at 370 nm. The OD of each sample dilution was calculated as the OD of the protein-coated wells minus the OD of the buffer-coated wells.

### Characterization of the ManBD (also known as ABS) of AcMBP and binding of anti-ManBD rAbs to trophozoites

To determine whether the ABS of AcMBP is the ManBD, an EcMBP-ABS fusion protein was made in the periplasm of *E. coli,* and its ability to bind to mannose-agarose resin (Sigma) and HCLE cells were determined. EcMBP-ABS was allowed to bind to mannose-agarose resin for one hour at 4°C in the binding buffer (0.1 M Tris buffer, pH 7.2, 100 mM NaCl and 20 mM CaCl_2_). The beads were washed three times, and the bound fraction was initially eluted using excess mannose (0.1 M α-Man) and then by boiling the beads in 2X SDS-PAGE sample buffer. EcMBP alone was a negative control, while ConA (Vector Laboratories) was a positive control. The samples were further analyzed by electrophoresis in 4-20% SDS-polyacrylamide gel.

HCLE cells, which were a gift from Vickery Trinkaus-Randall of Boston University Medical Center, were grown in Keratinocyte Serum-Free Medium (KSFM) with the following supplements: 25 µg/mL bovine pituitary extract, 0.02 nM EGF, 0.3 mM CaCl2, 100 U/ml Penicillin, and 100 µg/ml Streptomycin (Onochie 2019). HCLE cells were grown to confluence in 3 mm glass bottom culture discs, fixed with 4% paraformaldehyde at room temperature for 15 minutes, washed three times with 1X PBS, and blocked with 1% BSA for one hour at room temperature. HCLE cells were stained with 10 ug of the EcMBP-ABS fusion-protein conjugated with AF594 along with anti-alpha tubulin antibody (Sigma) conjugated with AF488 for 1 hour at room temperature. HCLE cells were also incubated with EcMBP alone conjugated with AF594 (negative control) or ConA conjugated with AF647 (positive control. Subsequently, the HCLE cells were washed three times with 1X PBS, mounted in VECTASHIELD® Antifade Mounting Medium (Vector Laboratories, Newark, CA), and visualized using 488 nm (Alexa flour 488) and 594 nm (Alexa flour 594) laser excitation.

Rabbit antibodies to EcMBP-ABS, which are referred to as anti-ManBD rAbs, were made, purified, and labeled with AF488, as described above for rAbs to cyst wall proteins. Dividing trophozoites from 11 isolates of *Acanthamoeba*, which were used to test the binding to CFW-labeled cysts by rAbs to wall proteins, were collected by centrifugation, washed three times with PBS, and fixed in 4% paraformaldehyde for 15 minutes at room temperature. The cells were labeled with anti-ManBD rAbs, labeled with DAPI, and visualized with high-power confocal microscopy, as described above. In three separate experiments, low-power images of 100+ trophozoites were captured and counted to determine the efficiency of detection of DAPI-labeled trophozoites of each isolate. Finally, anti-ManBD rAbs bound to trophozoites of the Neff strain of Ac were visualized with a conventional Zeiss Axio Observer Z1 microscope.

## Abbreviations

4DK: four disulfide knots
ABS: antiparallel β-sandwich
Ac: *Acanthamoeba castellanii*
AcMBP: Ac mannose-binding protein
AK: *Acanthamoeba* keratitis
ATCC: American Type Culture collection
BHF: β-helical fold
BJRF: β-jelly-roll fold
CFW: calcofluor-white
ConA: Concanavalin A
CRR: Cys-rich region
CuRO-1: first copper oxidase domain
EcMBP: *E. coli* maltose-binding protein
ELISA: enzyme-linked immunoassay
HCLE: human corneal limbal epithelial cell
ManBD: mannose-binding domain
rAb: rabbit antibodies
WGA: wheat germ agglutinin

## Acknowledgments

This work was supported in part by grants to JS from National Institutes of Health (R01 GM129324) and from Howard Hughes Medical Institute (Emerging Pathogens Initiative).

## Author contributions

Bharath Kanakapura Sundararaj and Manish Goyal each designed, performed, and analyzed the experiments, as well as prepared figures and wrote a draft of the paper. John Samuelson provided funding, supervised their work, and finished writing the paper.

