## Supplemental Figures for "Identification of new targets for the diagnosis of cysts (four) and trophozoites (one) of the eye pathogen *Acanthamoeba*"

Fig. S1. Sequences used for making *E. coli* maltose-binding protein-fusions for immunizing rabbits.

Fig. S2. Nucleotide sequences of 18S RNAs of 11 *Acanthamoeba* isolates from the ATCC.

Fig. S3. Low-power confocal micrographs of cysts labeled with calcofluor-white and rabbit antibodies to wall proteins.

Fig. S4. Images with a conventional fluorescence microscope of *Acanthamoeba* cysts and trophozoites labeled with rabbit antibodies.

Fig. S5. Disulfide bonds in the antiparallel β-sandwich of AcMBP-1 and Western blots of the native protein.

Fig. S6. Low power confocal micrographs of human corneal limbal epithelial cells and *Acanthamoeba* trophozoites.

### A. Jonah-1, Luke-2, Leo-A, laccase-1, and ManBD

```
>Jonah-1 (ACA1_164810)
VGSSCPAEDFFGPAADLNVLVTGSDKLIYDLDMVNGDVEGRVAVNGGFRVKSFGTAQAYSCPDTANYASMFNLIVNGKMDYSNGQLFCG
SSISLSSMNDPSLRQKDFGATNARVAAGKTIKDVTGFDFVATAGYLQGVSNYFSKYAPGTHSHFRVTEVSPADVNKLNIEGGDYVQVIN
IKGSGDINFSNFEIPTALNPSQVIYNVVGNNNIKISGFGLKGHLLALNSDVVLENGHVAGNVYVRSLVGQGSGQVNLAPCP

>Luke-2 (ACA1_377670)
QSCTASLNQTRQASWVDSEQFPRSLWTVEIRNTGQQAVTNVLLSIQGSINQIWEVVLVNDSLYRLPDWRLQVGGIPAGQSHVFGFIVNN
NSTAAPVSLVSVQCGNVTVPSPSPSASASSVPSPSASASAAPSPSAGASPSSSPRPSPSGGCSLLASQVARSGAGGSWSDNSNRFQIYD
ITLNNNGASRLTQASLTIAIAADQAIAQFWNLERVDATNTFNIESFLLPAPGGSQSGLGYVLQTPLNSTSDGSSIGAVNFRC

>Leo-A (ACA1_074730)
CPGSDFFPDQRGCCPVVVNSVPFYADAQGCFPRIISGVAFYADAGGCYPSSAGVYRSAGAVCPADRVCEPECPDMAIVVAEQQGGARLC
ANGFFSDVNGCCPRVLVDNVPSYRDAQGCYPISIASVIFYADVGRCYPNSSGVYRPAGAVCPAGQQCVVSCDRNYATS

>laccase-1 (ACA1_068450)
VKQTCDGDHYDITMGVVKHQFHPSLPVVDIYGYGGTYPGGTFEAVVNRPVHVTWRNSLPEKHILPTEPLEMEGMHMENPPDSAGVVHLH
GALIDPKDDGFSMDWITRGQKIQYHYTNEQLPLTMWYHDHVMTRTRVNVYAGLAGFYLLRDE

>ManBD of AcMBP-1 (AAT37864)
GTCNLSGAIKQPGLDCSSTSCSITSGTFPFPLPQGETYDSFYSWILGVIGTDGATVNAQYVDYTKADPNIYFTAGQTNCMVNLTFVYEV
AFYRNSMGYFTFTRDSKPTSVGSVTLKPVFSETTVDCSRTSSQPLPGTSCLAPGSTISLGPFSSTQAVGFYLKQDSICSGTTTFYSVDA
LNKVTSRWKPIPAAHGRMIAVLRDPNTLRAYLGFEDSPDGSDSDYNDNVFSVTSNCEIDTSLLP
```

### B. C-termini of CuRO-1 domains of laccases

```
Ac laccase-1   VKQTCDGDHYDITMGVVKHQFHPSLPVVDIYGYGGTYPGGTFEAVVNRPVHVTWRNSLPE    60
Ac laccase-2   YCTRRGVDKYVVTMSAIQHRLHPDLNLTQVYGYAGTYPGGTIEAVVDRPVAVTWRNHLPD    60
Ac laccase-3   VSRKCGVDRYEVTFNSFEQQLHPSLPPTTVYGYDGSYPGATFEAKVNRPVEVTWTNDLPL    60

Ac laccase-1   KHILPTEPLEMEGMHMENPPDSAGVVHLHGALIDPKDDGFSMDWITRGQKIQYHYTNEQL    120
Ac laccase-2   THFLPTEPLGHLG--SEPVPDSAAVPHLHGAHVPPDSDGDPMVWITRGQDADFYYPNGQL    118
Ac laccase-3   HHILPTVPLEG----SETPPESAGVVHLHGGHTPADSDGNPLDFIVKGQQQQFHYPNRQL    116

Ac laccase-1   PLTMWYHDHVMTRTRVNVYAGLAGFYLLRDE   151
Ac laccase-2   ATTMWYHDHIMGRTRVNVYAGLVGFYLLRDE   149
Ac laccase-3   ATTLWYHDHIWTRNRANVVAGLAGFYLLRDK   147
Clostridium    GTTMWYHDHGMGETRLNVYAGLAGFYILRDS   153
Bacillus       QAILWYHDHAMALTRLNVYAGLVGAYIIHD-   158
Bifiguratus    PATMWYHDHALGITRLNVYAGLVGFYLLR--   150
Daucus         PGNLWYHDHAMGLTRVNLLAGLIGSYIIR--   150
```

**Fig. S1. Sequences used for making *E. coli* maltose-binding protein-fusions for immunizing rabbits.** A. Jonah-1 contains a single β-helical fold, while Luke-2 contains two β-jelly-roll folds separated by Ser-rich spacer. Leo-A contains two sets of four disulfide knots, while laccase-1 contains the first copper oxidase domain. The mannose-binding domain (ManBD) of the *Acanthamoeba* mannose-binding protein (AcMBP-1, which is absent in AmoebaDB but present in NR database of NCBI, contains an antiparallel β-sandwich. B. Alignment marks in yellow identities with CuRO-1 domain of Ac laccase-1. Ac laccase-2 (ACA1_006180) had minor errors in protein prediction in AmoebaDB, which were corrected using an unpublished transcriptome and proteome made in collaboration with investigators at the Broad Institute. Ac laccase-3 (ACA1_008840) was severely truncated in AmoebaDB and so lacked the CuRO-1 domain, which was identified in the unpublished transcriptome and proteome. Other CuRO-1 sequences came from *Clostridium sp.* (SFU68967), *Bacillus cenocepacia* (4AK0), *Bifiguratus adelaidae* (OZJ01666), and *Daucus carota* (XP_017216893).

>30010

ATTTTCTGCCACCGAATACATTAGCATGGGATAATGGAATAGGACCCTGTCCTCCTATTTTCAGTTGGTTTTGGCAGCGCGAGGACTAG
GGTAATGATTAATAGGGATAGTTGGGGGCATTAATATTTAATTGTCAGAGGTGAAATTCTTGGATTTATGAAAGATTAACTTCTGCGAA
AGCATCTGCCAAGGATGTTTTCATTAATCAAGAACGAAAGTTAGGGGATCGAAGACGATCAGATACCGTCGTAGTCTTAACCATAAACG
ATGCCGACCAGCGATTAGGAGACGTTGAATACAAAACACCACCATCGGCGCGGTCGTCCTTGGCGTCTGTCCCTTTCAACGGGGGCAGG
CGCGAGGGCGGTTTAGCCCGGTGGCACCGGTGAATGACTCCCCTA

>50370
ATTTTCTGCCACCGAATACATTAGCATGGGATAATGGAATAGGACCCTGTCCTCCTATTTTCAGTTGGTTTTGGCAGCGCGAGGACTAG
GGTAATGATTAATAGGGATAGTTGGGGGCATTAATATTTAATTGTCAGAGGTGAAATTCTTGGATTTATGAAAGATTAACTTCTGCGAA
AGCATCTGCCAAGGATGTTTTCATTAATCAAGAACGAAAGTTAGGGGATCGAAGACGATCAGATACCGTCGTAGTCTTAACCATAAACG
ATGCCGACCAGCGATTAGGAGACGTTGAATACAAAACACCACCATCGGTGCGGTCGTCCTTGGCGTCTCGGTTTCGGCCGGGGTGCGGG
GACGGCTTAGCCCGGTGGCACCGGTGAATGACTCCCCTAGCAGCCTTGTGAGAA

>50324
TTCACCGGTGCCACCGGGCTAAGCCGCCCTCGCGCCGGCCCCGTGAAGGACCGACGCCAAGGACGACCGCGCCGATGGTGGTGTTTTGT
ATTCAACGTCTCCTAATCGCTGGTCGGCATCGTTTATGGTTAAGACTACGACGGTATCTGATCGTCTTCGATCCCCTAACTTTCGTTCT
TGATTAATGAAAACATCCTTGGCAGATGCTTTCGCAGAAGTTAATCTTTCATAAATCCAAGAATTTCACCTCTGACAATTAAATATTAA
TGCCCCCAACTATCCCTATTAATCATTACCCTAGTCCTCGCGCTGCCAAAACCAACTGAAAATAGGAGGACAGGGTCCTATTCCATTAT
CCCATGCTAATGTATTCGGTGGCAGAAAATTGGATCTGCCTGCTTTGAACACTCTAATTTT

>MEI_0184
TTCACCGGTGCCACCGGGCTAAGCCACCCCCGCGCTGGCTTTTGAAGACCAACGCCAAGGACGACCGCACCGATGGTGGTGTTTTGTAT
TCAACGTCTCCTAATCGCTGGTCGGCATCGTTTATGGTTAAGACTACGACGGTATCTGATCGTCTTCGATCCCCTAACTTTCGTTCTTG
ATTAATGAAAACATCCTTGGCAGATGCTTTCGCAGAAGTTAATCTTTCATAAATCCAAGAATTTCACCTCTGACAATTAAATATTAATG
CCCCCAACTATCCCTATTAATCATTACCCTAGTCCTCGCGCTGCCAAAACCAACTGAAAATAGGAGGACAGGGTCCTATTCCATTATCC
CATGCTAATGTATTCGGTGGCAGAAAATTGGATCTGCCTGCTTTGAACACTCTAATTTT

>30872
GATTCATTTTCTGCCACCGAATACATTAGCATGGGATAATGGAATAGGACCCTGTCCTCCTCTTTTCAGTTGGTTAATTTATGTGCGAG
GATCAGGGTAATGATTAATAGGGATAGTTGGGGGCATTAATATTTAATTGTCAGAGGTGAAATTCTTGGATTTATGAAAGATTAACTTC
TGCGAAAGCATCTGCCAAGGATGTTTTCATTAATCAAGAACGAAAGTTAGGGGATCGAAGACGATCAGATACCGTCGTAGTCTTAACCA
TAAACGATGCCGACCAGCGATTAGGAGACGTTGAATACAAAACACCGCTAAAGATAATTCATTATATGGCTTCACGGCTGTATAGTGTT
TGTCTTTAGTTCATGGTGAATGACTCCCCTAGC

>30137
GAGTCATTCACCGGTGCCACCGGcCTCCCCCGTCCCCGCACCCCGGCAGAAACCGAGACGCCAAGGACGACCGCACCGATGGTGGTGTT
TTGTATTCAACGTCTCCTAATCGCTGGTCGGCATCGTTTATGGTTAAGACTACGACGGTATCTGATCGTCTTCGATCCCCTAACTTTCG
TTCTTGATTAATGAAAACATCCTTGGCAGATGCTTTCGCAGAAGTTAATCTTTCATAAATCCAAGAATTTCACCTCTGACAATTAAATA
TTAATGCCCCCAACTATCCCTATTAATCATTACCCTAGTCCTCGCGCTGCCAAAACCAACTGAAAATAGGAGGACAGGGTCCTATTCCA
TTATCCCATGCTAATGTATTCGGTGGCAG

>PRA115
ATTTTCTGCCACCGAATACATTAGCATGGGATAATGGAATAGGACCCTGTCCTCCTATTTTCAGTTGGTTTTGGCACGCGAGGACCAGG
GTAATGATTAATAGGGATAGTTGGGGGCATTAATATTTAATTGTCAGAGGTGAAATTCTTGGATTTATGAAAGATTAACTTCTGCGAAA
GCATCTGCCAAGGATGTTTTCATTAATCAAGAACGAAAGTTAGGGGATCGAAGACGATCAGATACCGTCGTAGTCTTAACCATAAACGA
TGCCGACCAGCGATTAGGAGACGTTGAATACAAAACACCACCATCGGTGCGGTCGTCCTTGGCGCGTCGCGGCTTGCCGCGGCGTGCGA
GGGCGGTTTAGCCTGATGGCATCGGTGAATGACTCCCC

>PRA411
ACTTTACCTGGGATATATGGAATTAGAACCCGGCCTCTTATTTTTCNTNGTTTTGTGTGTTCAATGCCTGCGCTAACGATTGGCTGGCA
TATGTGGGGGCATTAATATTTAATTGTGGGAAGTGATATTCTTGGATTTTTGGGGGATTAACTTTTGCATTTTCATCTGTCAAGTTTGT
TTTTTTTAATCAAGAAAGATTGTTANGGGATCGAAGACAAGCAGATTCCGTTTNANTCTTAACCAAAATCGAGGCCGACAAGCGATTAA

GATACGTTCAAAACCTAACCCCAACACCAGTCCGGCCGTCCTTGGCGTCACGTTTCCNNCCGGGGTGCGGGGGGGGGGGGCCCCGCGG
TTCCCCCCAATGAAACCGCCAGCANNNTGCGNCAATCCTTTGGGCGGCTCCCCCTAAAGAATAACCCCCCTACC

>50655
TTCACCGGGCCATCAGGGGAATGAGACCGCCTCCACTTCCCCGTTGCCGGAAAAGCGAAGCACGACCCCACTGCCTGATGGTGGTGTTT
TGTATTCAACGTCTCCTAATCGCTGGTCGGCATCGTTTATGGTTAAGACTACGACGGTATCTGATCGTCTTCGATCCCCTAACTTTCGT
TCTTGATTAATGAAAACATCCTTGGCAGATGCTTTCGCAGAAGTTAATCTTTCATAAATCCAAGAATTTCACCTCTGACAATTAAATAT
TAATGCCCCCAACTATCCCTATTAATCATTACCCTGGTCCTCAAAAACCAACGCGGAAAATAGGAGGACAGGGTCCTATTCCATTATCC
CATGCTAATGTATTCGGTGGCAGAATGGTAAAATCTGCCTGCTTTGAACACTCTAATTTT

>ESBC4
TTCACCGGTGCCACCGGGCTAAGCCGCCCTCGCGCCGGCCCCGTGAAGGACCGACGCCAAGGACGACCGCGCCGATGGTGGTGTTTTGT
ATTCAACGTCTCCTAATCGCTGGTCGGCATCGTTTATGGTTAAGACTACGACGGTATCTGATCGTCTTCGATCCCCTAACTTTCGTTCT
TGATTAATGAAAACATCCTTGGCAGATGCTTTCGCAGAAGTTAATCTTTCATAAATCCAAGAATTTCACCTCTGACAATTAAATATTAA
TGCCCCCAACTATCCCTATTAATCATTACCCTAGTCCTCGCGCTGCCAAAACCAACTGAAAATAGGAGGACAGGGTCCTATTCCATTAT
CCCATGCTAATGTATTCGGTGGCAGAAAATTGGATCTGCCTGCTTTGAACACTCTAATTTT

>50676
GATCCATTTTCTGCCACCGAATACATTAGCATGGGATAATGGAATAGGACCCTGTCCTCCTATTTTCAGTTGGTTTTGGCAGCGCGAGG
ACTAGGGTAATGATTAATAGGGATAGTTGGGGGCATTAATATTTAATTGTCAGAGGTGAAATTCTTGGATTTATGAAAGATTAACTTCT
GCGAAAGCATCTGCCAAGGATGTTTTCATTAATCAAGAACGAAAGTTAGGGGATCGAAGACGATCAGATACCGTCGTAGTCTTAACCAT
AAACGATGCCGACCAGCGATTAGGAGACGTTGAATACAAAACACCACCATCGGCGCGGTCGTCCTTGGCGTTGTCGGCTTCACGGCTGG
CGGCGCGAGGGCGGTTTAGCCCGGTGGCACCGGTGAATGACTCCCC

**Fig. S2. Nucleotide sequences of 18S RNAs of 11 *Acanthamoeba* isolates from the ATCC.** Each of these sequences were compared with those used to identify and tree genotypes of *Acanthamoeba*.

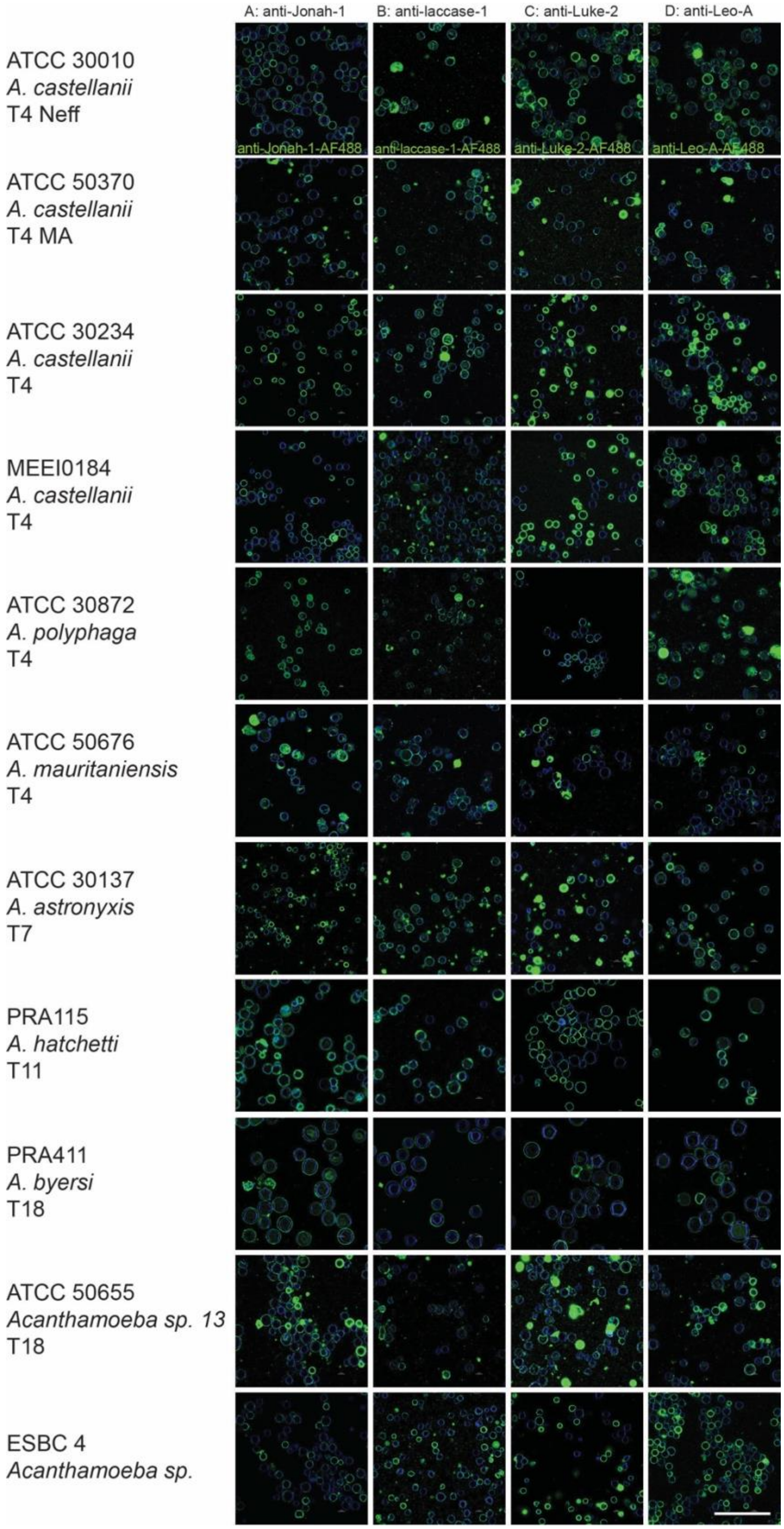


**Fig. S3. Low power confocal micrographs of cysts labeled with calcofluor-white and rabbit antibodies to wall proteins**. A to D. Cysts of 11 *Acanthamoeba* isolates (shown at high power in Fig. 2 in the text) were labeled with Protein A-purified antibodies tagged with Alexa Flour 488, and the percentage of CFW-tagged cysts detected was counted for two experiments and plotted in Fig. 3A in the text. A single scale bar for A to D is 50 μm.

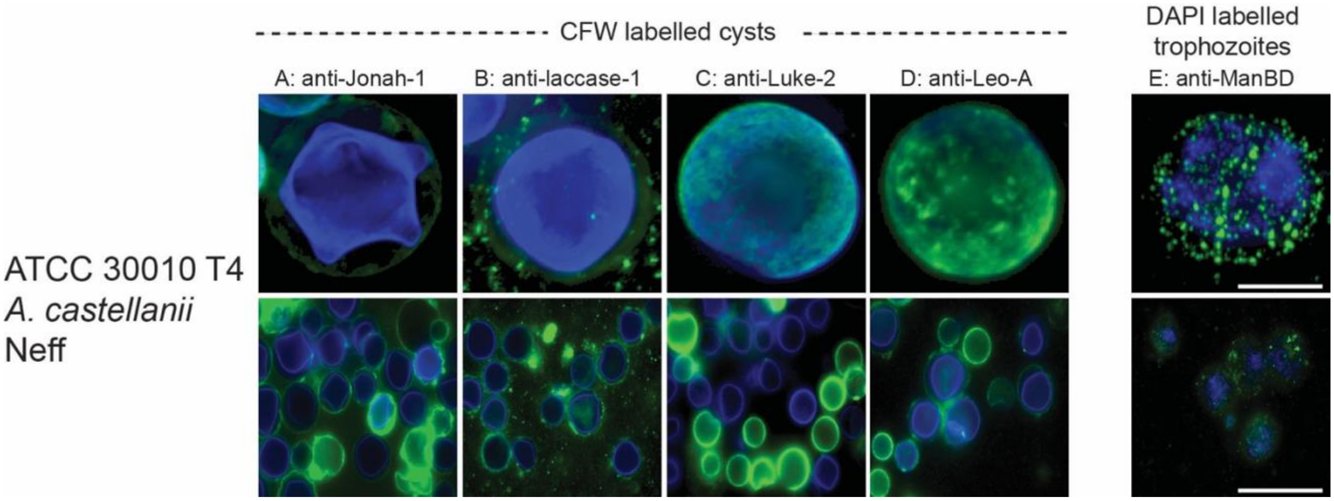


**Fig. S4. Images with a conventional fluorescence microscope of *Acanthamoeba* cysts and trophozoites labeled with rabbit antibodies.** A to D. High power (top row) and low power (bottom row) micrographs of rabbit antibodies to wall proteins binding to calcofluor-white labeled cysts. E. High (top) and low (bottom) power micrographs of rabbit antibodies to the mannose-binding domain binding to DAPI-stained trophozoites. Scale bar for top row is 5 µm and for bottom row is 50 µm.

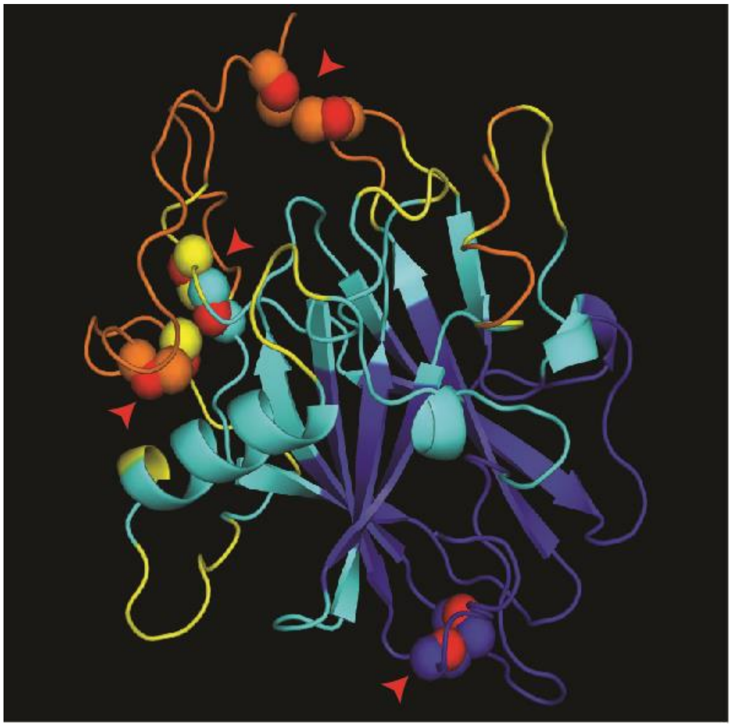

**Fig. S5. Disulfide bonds in the antiparallel β-sandwich (ABS) of AcMBP-1.** The ABS of AcMBP-1 (see also Figs. 4A to 4C in the text) contains four disulfide knots (red arrowheads), which connect peripheral loops.

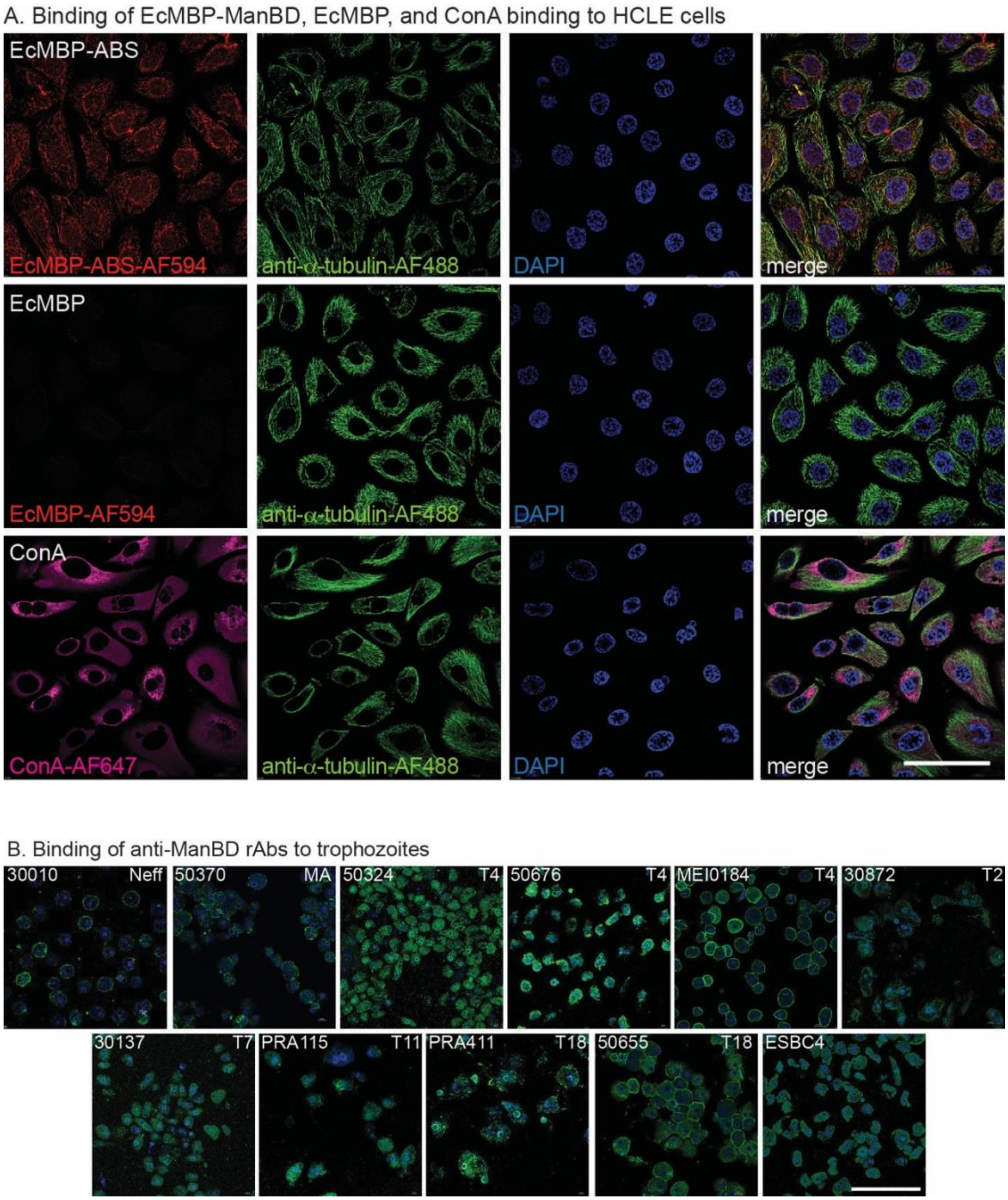


**Fig. S6. Low power confocal micrographs of human corneal limbal epithelial cells and *Acanthamoeba* trophozoites.** A. An EcMBP-fusion to the ABS of AcMBP-1, which is also called ManBD, binds to HCLE cells labeled with anti-tubulin antibodies and DAPI, while EcMBP alone (negative control) does not bind. The plant lectin Concanavalin A (positive control) also binds to HCLE cells. High power confocal micrographs of HCLE cells labeled with EcMBP-ABS and ConA are show in figs 4G and 4H in the text. B. Trophozoites of 11 *Acanthamoeba* isolates (shown at high power in Fig. 4I in the text) were labeled with Protein A-purified antibodies tagged with Alexa Flour 488, and the percentage of trophozoites detected was counted for two experiments and plotted in Fig. 4J in the text. Scale bar for A and B are each 50 μm.
